# Fucoidan carbon is stored in coastal vegetated ecosystems

**DOI:** 10.1101/2024.12.02.624615

**Authors:** Inga Hellige, Aman Akeerath Mundanatt, Jana Massing, Jan-Hendrik Hehemann

## Abstract

Coastal vegetated ecosystems are key-nature based solutions for climate change mitigation. Mangroves, seagrass meadows and saltmarshes contribute to carbon sequestration not only through their photosynthetic activity but also by anchoring sediments with their extensive root systems. By modulating flow coastal vegetation creates a low energy environment for sediment that includes carbon to accumulate. These roots physically stabilize the sediment, prevent erosion and enhance long-term retention of organic carbon. Hence, we hypothesized marine, algae derived organic matter may especially accumulate in plant vegetated ecosystems. We used algal and plant glycans as carbon sequestration proxy to trace the input and stabilization from source to sink and found those molecules in 93 sediment cores across different coastal vegetated ecosystems from temperate to tropical regions. Specific monoclonal antibodies showed algal-derived fucoidans were present in sediments of coastal vegetated ecosystems. Our findings suggest that the restoration of plant ecosystems that fix carbon dioxide, protect coasts and enhance biodiversity should also be enumerated for the stored carbon from distant donors. Conclusively, carbon sequestration is a synergistic outcome of photosynthetic contributors acting in concert across different ecosystems.

## 1 Introduction

Coastal vegetated ecosystems, including mangroves, seagrass meadows, and saltmarshes, play a crucial role as carbon sinks, storing estimated amounts of up to 0.7 Pg carbon year^-1^ (Temmink et al., 2022). With carbon sequestration rates up to 10 times greater than those of terrestrial forests (Mcleod et al., 2011), these ecosystems are increasingly recognized as nature-based climate solutions. This high carbon sequestration capacity, coupled with their abilities to protect coastlines, support biodiversity, and improve water quality, underscores their significance in carbon dioxide removal (CDR) strategies (Gattuso et al., 2018; Hagger et al., 2022; Mengis et al., 2023).

Mangroves, seagrass and saltmarshes sequester carbon through biomass and sediment storage and by releasing dissolved organic carbon (DOC) through their root system. Seagrass meadows were shown to be enriched in the sugar sucrose compared to unvegetated areas (Sogin et al., 2022). In contrast, macro- and microalgae, which lack root systems, release most of their carbon as exudates, constituting between 1% and 35% of their net primary production (Abdullah and Fredriksen, 2004; Paine et al., 2021; Reed et al., 2015; Wada et al., 2007; Wada and Hama, 2013). Due to the challenges in tracking algae’s carbon storage potential from source to sink, they are often excluded from blue carbon strategies (Krause-Jensen et al., 2018; Krause-Jensen and Duarte, 2016).

Despite these challenges, algae contribute substantially to blue carbon pools. Macro- and microalgae have been shown to supply up to 50% carbon in seagrass sediments (Kennedy et al., 2010) and up to 60% carbon of Red Sea mangrove sediments (Almahasheer et al., 2017), identifying algae as significant carbon donors (Krause-Jensen et al., 2018; Ortega et al., 2019). However, once released into the environment, tracking algae-derived carbon becomes complex. While a portion of this carbon is remineralized, 10-60% can persist under environmental conditions, potentially forming long-term carbon sinks (Filbee-Dexter and Wernberg, 2020; Krause-Jensen and Duarte, 2016; Wada et al., 2008; Watanabe et al., 2020).

Further complicating this tracking process is the complexity of algae exudates. For instance, brown algae and diatoms secrete the complex and variable extracellular matrix polysaccharide fucoidan that might resist microbial degradation (Arnosti et al., 2012; Buck-Wiese et al., 2023; Giljan et al., 2023; Lloyd et al., 2022; Vidal-Melgosa et al., 2021). These substances can assemble into particles (Huang et al., 2021; Vidal-Melgosa et al., 2021) that are carried by tides and currents before sinking to sediment (Salmeán et al., 2022; Vidal-Melgosa et al., 2022).

The extent of carbon transport and the composition of the stored organic carbon in coastal vegetated ecosystems remains understudied. Tracing carbon from source to sink is essential for understanding the potential for long-term carbon storage. Here, we hypothesized that polysaccharides, such as fucoidan can serve as tracers for algal carbon stored within coastal vegetated ecosystems, suggesting that blue carbon sequestration results from the collective carbon sequestration across ecological communities.

## 2 Methods

### Sampling

50cm deep sediment cores were collected with a Russian peat corer (5cm diameter) in saltmarsh, seagrass, mangroves and unvegetated areas around the German Bight, Malaysia and Columbia (**Fig. S1A-C**). Up to 3 points per ecosystem were sampled along a transect (**Fig. S1D**), in total 93 cores were analysed (**Table S1**). Additionally, porewater samples were taken using lances or digging of holes. Samples were taken between 30 and 50 cm along 9 points per ecosystem at North Sea sites (**Table S2**). Samples were directly filtered over pre-combusted (450°C, 4.5h) GMF and GF/F filters and 200 ml of filtered porewater samples were frozen at -20°C for polysaccharide analysis.

### Processing

Cores were split visually according to physiochemical layers into up to 5 parts. Each layer was homogenized by hand in a ziplock bag and stored at 4°C. Subsamples were frozen at -20°C prior to freeze drying with an Alpha 1-4 LSCbasic freeze dryer from Christ at -55°C until constant vacuum. Samples were pulverised with a ball mill (FRITSCH LLC Planeten-Mikromühle PULVERISETTE 7 premium line) at 500 rpm for 3 min. Sediment powder was stored at room temperature in the dark until further analysis.

Porewater samples were run over AMICON filtration device with 5kDa membrane (Biomax) to separate the polysaccharide fraction over 5 kDa, washed 2 times with 300 µL MilliQ and up-concentrated by a factor of 5 in MilliQ-water. Samples were freeze dried and resuspended at a final concentration factor of 100 times in MilliQ-water.

### Carbohydrate extraction

90 mg of dried and pulverised material was sequentially extracted using MilliQ-water and 0.3 M EDTA. Sediment was mixed with 1.8 mL of MilliQ-water, vortexed and kept in an ultrasonic water bath for 1 hour. Extracts were centrifuged at 6000G for 15 min. The resulting supernatant was transferred into a new vial and the pellets were mixed with 1.8 mL of 0.3 M EDTA, vortexed and kept in an ultrasonic water bath for 1 hour. Extracts were centrifuged at 6000G for 15 min and the supernatant was again transferred into a new vial. Extracts were frozen at -20° until further analysis.

### Phenol-sulfuric assay

The total carbohydrate content was determined based on (Dubois et al., 1951). In short, 100 µL of resuspended samples or extracts were mixed with 100 µL of 5% phenol solution and 500 µL of concentrated sulfuric acid, then incubate for 10 minutes at RT and afterwards for 20 min at 30°C. Absorbance at 490 nm was measured using Spectramax Id3 plate reader (Molecular Devices) against a glucose standard curve.

### Acid hydrolysis

Polysaccharides were acid hydrolysed into quantifiable monosaccharides by adding 300 µL 2M HCl to 300 µL of the porewater extracts and MilliQ-sediment extracts and the 20x diluted EDTA sediment extract in pre-combusted glass vials (450°C, 4.5 h). Glass vials were sealed and polysaccharides hydrolysed at 100°C for 24 h. The samples were transferred after hydrolysis to microtubes and dried in an acid-resistant vacuum concentrator (Martin Christ Gefriertrocknungsanlagen GmbH, Germany). Samples were resuspended in 300 µL MilliQ-water.

### Monosaccharide quantification

As previously described in two studies (Engel and Händel, 2011; Vidal-Melgosa et al., 2021), monosaccharides were quantified using anion exchange chromatography with pulsed amperometric detection (HPAEC-PAD). The sample analysis was conducted on a Dionex ICS-5000+ system, equipped with a CarboPac PA10 analytical column (2 x 250 mm) and a CarboPac PA10 guard column (2 x 50 mm). Neutral and amino sugars were separated applying an isocratic phase using 18 mM NaOH. A gradient reaching up to 200 mM NaCH3COO was applied to separate acidic monosaccharides.

### Microarray

MilliQ-water and EDTA sediment extracts were equally combined, 30 µL were transferred into wells of 384-microwell plates, where two consecutive two-fold dilutions were done in printing buffer (55.2% glycerol, 44% water, 0.8% Triton X-100). The microwell plates were centrifuged at 3500 x g for 10 min at 15 °C. Microarray printing and probing was performed as previously described in a recent study (Vidal-Melgosa et al., 2022).

### ELISA

Polysaccharide were detected using enzyme-linked immunosorbent assay (ELISA) as described in the following studies (Cornuault et al., 2014; Vidal-Melgosa et al., 2021). In short, 100 µl of each combined sediment extract and the porewater samples were pipetted into a pre-coated 96 well plate and incubated overnight at 4°C. Signal was developed using primary antibodies (BAM1 for porewaters samples and BAM1 and JIM13 for sediment samples) in skim milk PBS solution at a concentration of 1:10, followed by antibody anti-rat in skim milk PBS solution at a concentration of 1:1000. Absorbance after development was measured at 450 nm using Spectramax Id3 plate reader (Molecular Devices).

### Statistical analysis

All statistical analysis were carried out using R4.4.1 (R core team (Anon, 2020), and Julia (Bezanson et al., 2017). The relative abundance of each monosaccharide was determined, all samples with a relative abundance of acidic sugars of over 50% were excluded from calculations. Pairwise distances were calculated using the Bray-Curtis distance metric for non-metric multidimensional scaling, using the vegan package (Oksanen et al., 2001). As NMDS can be quite sensitive to (Fahimipour and Gross, 2020), we also applied diffusion maps (Coifman et al., 2005) as a nonlinear dimensionality reduction method for comparison using 1/Bray-Curtis distance to construct the similarity matrix and a threshold of 5. The eigenvectors corresponding to the smallest non-zero eigenvalues are the most interesting as they identify the directions of the largest variation, i.e. the main dimensions of the data manifolds.

To test for significant changes between ecosystems combining MiliQ-water and EDTA extracts we applied the following statistical methods using the car package (Pante and Simon-Bouhet, 2013). In case of no normal distribution, we performed the Kruskal-Wallis test, followed by the pairwise Wilcoxon test with α-correction according to Bonferroni to adjust for the inflation of type I error due to multiple testing. Bonferroni adjustment was performed as follows: p= 0.05/k, with k as the number of single hypotheses. Here, k= 6 was used for comparison among all 4 ecosystems and k= 3 for locations without mangroves. Therefore, α= 0.0083 or α= 0.0167 was considered statistically significant. When normality was met, we conducted an ANOVA, followed by a post-hoc Tukey test for pairwise comparisons.

## 3 Results

### 3.1 Global sediment analysis revealed similar monosaccharide abundance despite different ecosystems and locations

To investigate the mono- and polysaccharides stored in sediments across coastal vegetated ecosystems, we analysed 93 sediment cores from the North Sea, Baltic Sea, Malaysia and Columbia. 50cm deep sediment cores were taken in saltmarsh, seagrass, mangroves and unvegetated areas. Up to 3 points per ecosystem were sampled along a transect (**Fig. S1, Table S1**). The major building blocks of polysaccharides, including fucose, galactose, glucose, mannose, xylose and galacturonic acid were found to be shared between all ecosystems and depths. Applying a non-metric multidimensional scaling (NMDS) analysis to the dataset of relative abundances of monosaccharides in each sample, we found no distinct clusters among the 93 sediment cores, indicating the composition of each sample remains unchanged with location and ecosystem (**Fig. 1A**). Diffusion mapping the dataset of relative abundances, revealed a reverse relationship between fucose and glucose, where high fucose abundances occur with low glucose abundances and vice versa (**Fig. 1B-C**).

**Figure 1:**
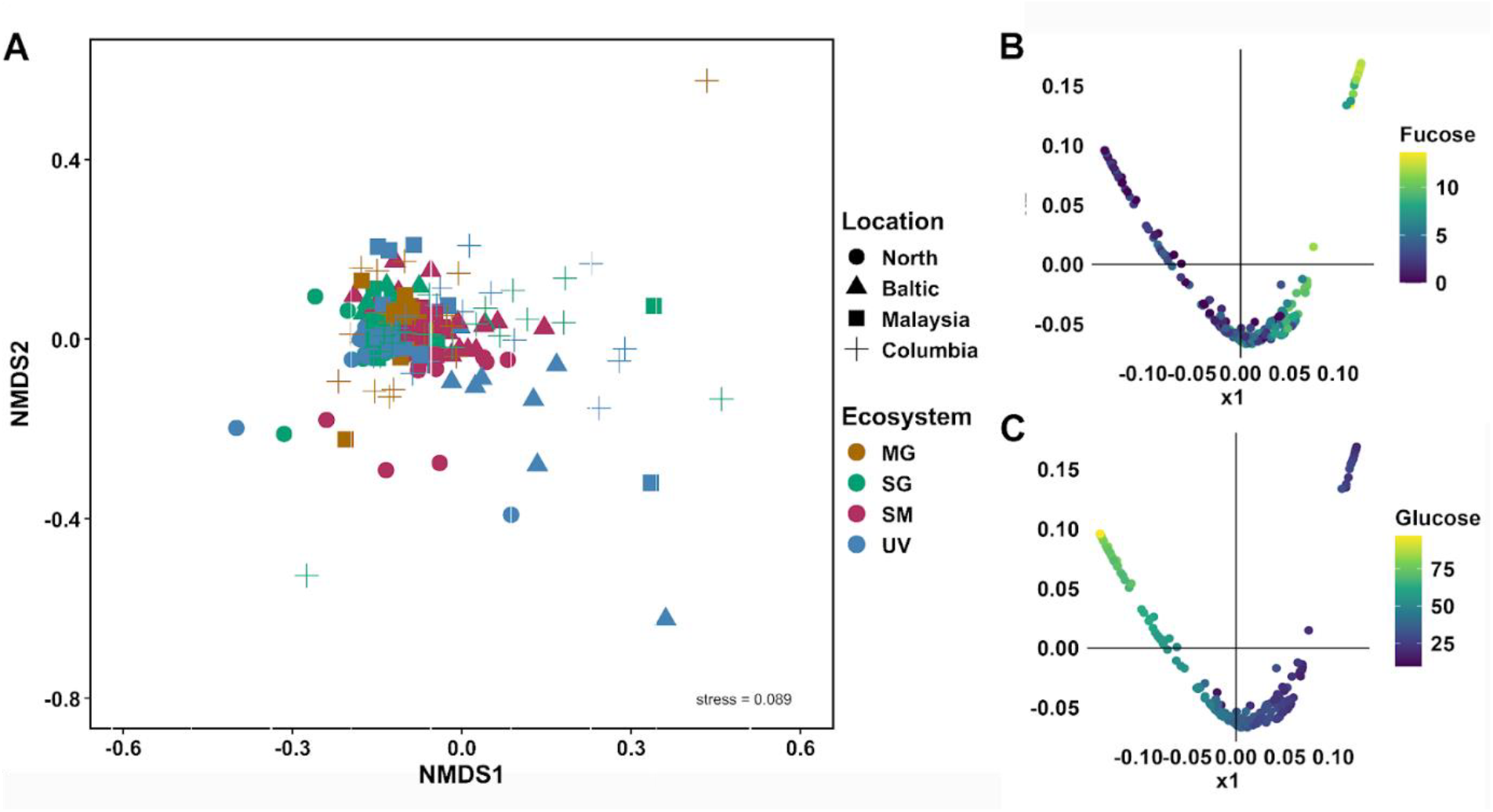
93 analysed sediment cores from temperate to tropical regions revealed no major differences in monosaccharide abundances, despite diverse ecosystems and locations. **A)** Non-metric multidimensional scaling (NMDS) plot, calculated by Bray-Curtis dissimilarity of monosaccharide composition of MilliQ-water and EDTA extracts across different sediment layers for four locations, North Sea and Baltic Sea in Germany, Malaysia and Columbia for four ecosystems, mangroves (MG, brown), seagrass (SG, green), saltmarsh (SM, maroon) and unvegetated (UV, blue). NMDS stress level of 0.089. **B-C)** Diffusion maps of the relative abundances of monosaccharides identify new explanatory variables. X1 and x2 correspond to two new variables that identify the main dimensions of the data manifolds. Colouring the datapoints by relative abundance of fucose **(B)** and glucose **(C)** reveals an inverse relationship, where samples with high fucose levels tend to have low glucose levels, and vice versa.

The quantification of total carbohydrates across different ecosystems revealed the highest maximum mean concentrations of 1.67 ± 0.23 mg g_dw_^-1^ in saltmarsh sediments. Mangroves and saltmarshes showed significantly higher concentrations compared to seagrass and unvegetated areas (p<0.001) (**Fig. 2A**). Of the total carbohydrates, 10% could be attributed to total hydrolyzable carbohydrates. Except for mangroves, all ecosystems showed significant differences to each other (p<0.001, after Bonferroni correction, α=0.0083) (**Fig. 2B**). Unvegetated areas ranged the lowest in mean concentrations of 35.94 ± 1.04 ug g_dw_^-1^, followed by seagrass with 67.65 ± 2.32 ug g_dw_^-1^ and saltmarsh (250.55 ± 9.04 ug g_dw_^-1^) areas. Mangroves recorded the highest mean concentrations of 494.08 ± 45.51 ug g_dw_^-1^, but also the greatest variations (**Fig. 2B**).

**Figure 2:**
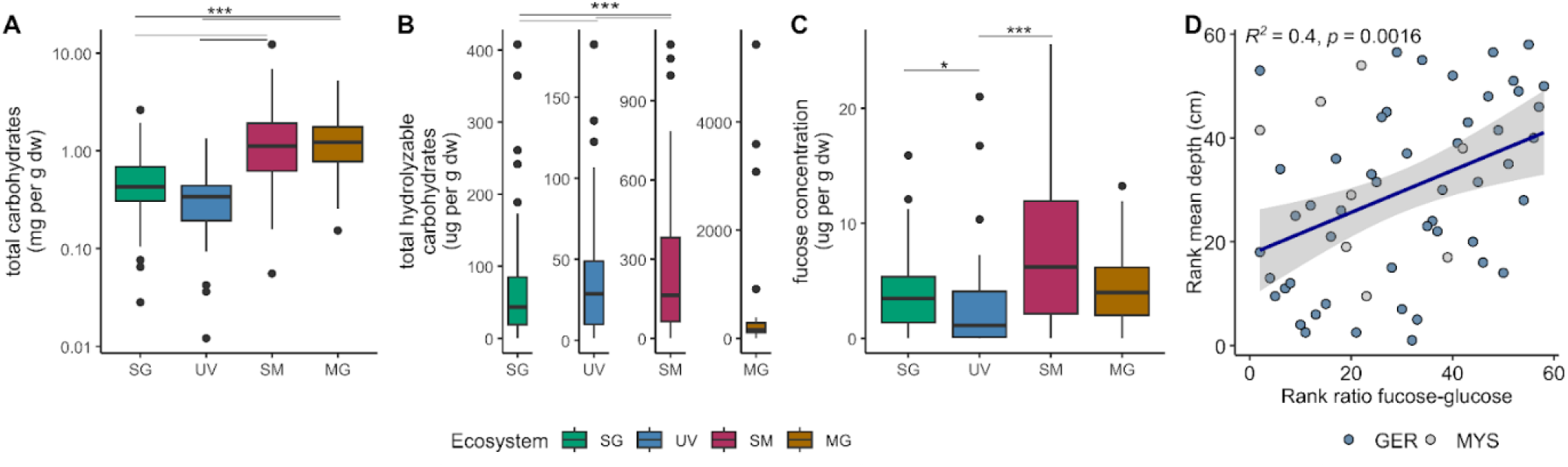
Up to 10% of total hydrolyzable carbohydrates can be attributed to fucose. **A)** Total carbohydrates, **B)** total hydrolyzable carbohydrates, **C)** fucose concentration in mg per g dry sediment from seagrass (SG; green), unvegetated (UV; blue), saltmarsh (SM; maroon) and mangroves (MG; yellow). **D)** Spearman rank correlation of mean depth in cm to fucose-glucose ratio (Germany: blue; Malaysia: lightgrey) indicating increase in fucose over glucose with sediment depths.

Fucose concentrations averaged 4.60 ± 0.34 ug g_dw_^-1^ across all samples (**Fig. 2C**). Significant differences were observed among concentrations between seagrass and unvegetated (p= 0.0066) and saltmarsh and unvegetated sediments (p<0.001, after Bonferroni correction, α = 0.0083). Mean fucose concentrations were lowest in unvegetated areas with 2.57 ± 0.46 ug g_dw_^-1^, followed by seagrass with 3.88 ± 0.40 ug g_dw_^-1^, by mangroves with 4.35 ± 0.67 and highest in saltmarshes with 7.79 ± 0.89 ug g_dw_^-1^ (**Fig. 2C**). Spearman rank correlation of mean depth to fucose-glucose ratio revealed an increase in fucose to glucose ratio with increasing sediment depth (R^2^= 0.4, p=0.0016; **Fig. 2D**).

### 3.2 Specific antibody binding shows algae glycans in sediment cores of coastal vegetated ecosystems

Structure specific monoclonal antibodies analysis imply algae as a source of glycans in coastal vegetated ecosystems. Analysis of carbohydrate microarrays indicates the presence of alginate, pectin, fucoidan, arabinogalactan protein glycan, (1→3)-β-D-glucan and grass-xylan (**Fig. S2**). Most signals were revealed for BAM7, indicative for alginate, fucoidan or pectin, across all ecosystems in Columbia and Germany. Further JIM13 indicative for arabinogalactan-protein glycans showed signal in saltmarsh and mangrove ecosystems. Notably, the upper layer of saltmarsh cores (0 up to 9.5 cm) in Heiligenhafen (HEI), Baltic Sea, revealed signal for grass xylan (LM27), not detected in further samples (**Fig. S2**). To confirm the presence of arabinogalactans and fucoidans the more sensitive antibody method ELISA was performed. Antibody binding to arabinogalactan-protein glycan (JIM13) confirmed the signal in saltmarshes and mangroves, a lower signal in seagrass sediments and close to no signal in unvegetated areas (**Fig. S3**). Monoclonal antibody BAM1, specific to algal polysaccharide fucoidan showed signal for all coastal vegetated ecosystems in all locations (**Fig. 3A**). Relative signals appeared to be stronger in coastal vegetated ecosystems, enriched in sediments of saltmarshes, mangroves and seagrass, slightly lower in unvegetated areas. Fucoidan stored in sediments of all coastal ecosystems was supported by the fucoidan antibody signal positively correlating with fucose concentration (p<0.0001 for seagrass, saltmarsh and unvegetated areas; mangroves: p= 0.0022) (**Fig. 3B**).

**Figure 3:**
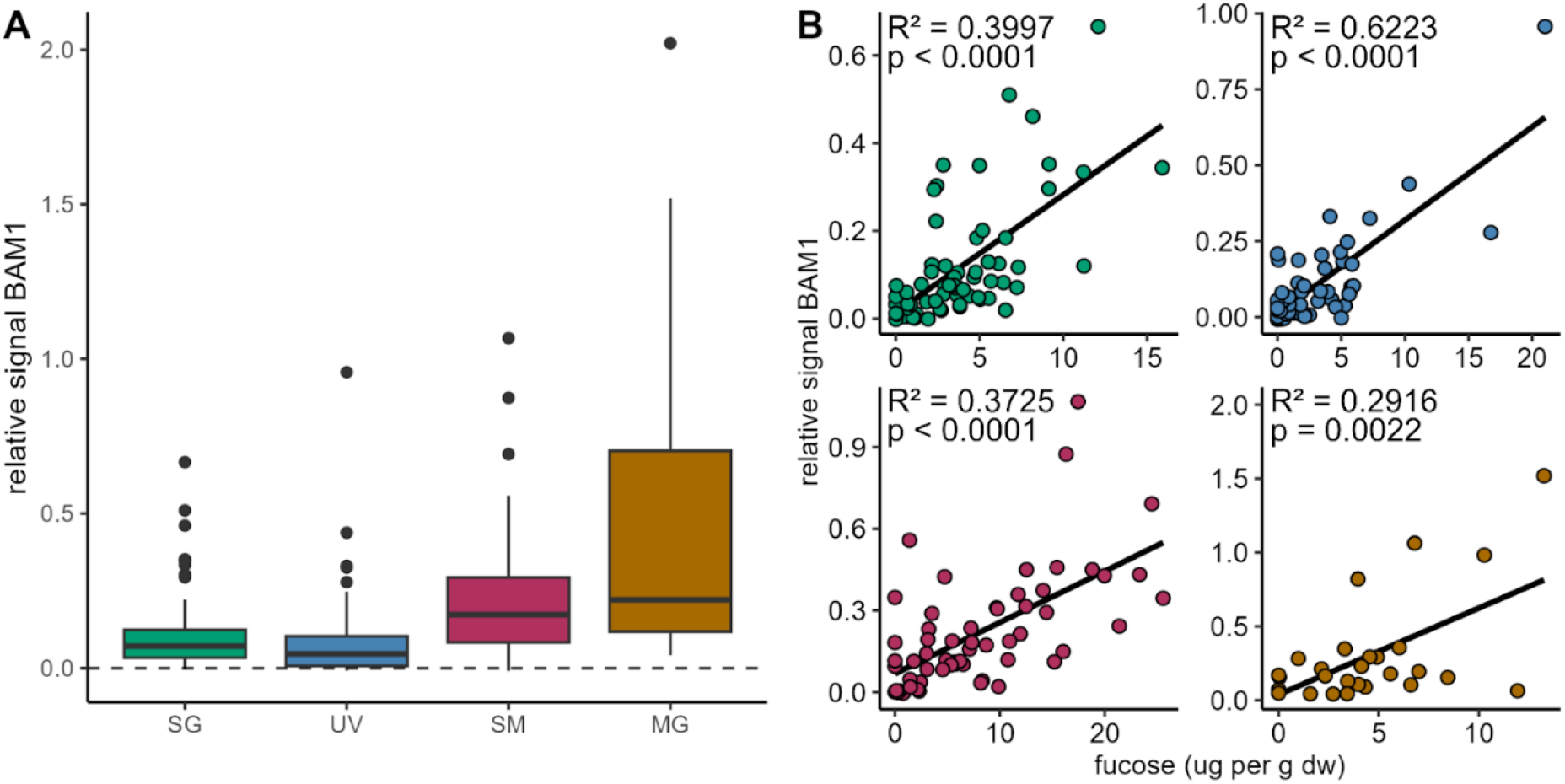
Algal glycan fucoidan correlates with fucose concentrations in sediments. **A)** Relative signal for antibody BAM1 (indicating presence of fucoidan) binding to sediment extracts from seagrass (SG; green), unvegetated (UV; blue), saltmarsh (SM; maroon) and mangrove areas (MG; brown). Relative signal normalized to MilliQ-water blank shown by dashed line. **B)** Relative BAM1 signal correlates with fucose concentration derived from monosaccharide analysis in ug g_dw_^-1^ (dw: dry weight).

The presence of the algal-derived glycan fucoidan was confirmed in porewater samples taken at depths of 30 to 50 cm along transects in North Sea sites (**Table S2**), using BAM1 antibody binding. The relative signal for fucoidan was notably higher beneath seagrass meadows compared to unvegetated areas. Similarly, strong signals were detected within the pioneer saltmarsh zone, relatively decreasing within the low saltmarsh zone and being absent inside the high saltmarsh (**Fig. 4A**). These findings align with measured fucose concentrations of 46.98 ± 26.76 µg L^-1^ within the pioneer saltmarsh zone and 40.52 ± 10.85 µg L^-1^ beneath seagrass meadows. Concentrations were lower in the low saltmarsh zone (26.77 ± 11.72µg L^-1^) and unvegetated areas (14.56 ± 2.78 µg L^-1^), with no detectable fucose in the high saltmarsh zone (**Fig. 4A**). These results indicate that dissolved algal-derived glycans, such as fucoidan, are secreted and transported into vegetated coastal ecosystems, which may act as acceptor systems. Algal-derived glycans carried by tidal waters may contribute to the enhancement and stabilization of sediment build-up within these ecosystems (**Fig. 4B**).

**Figure 4:**
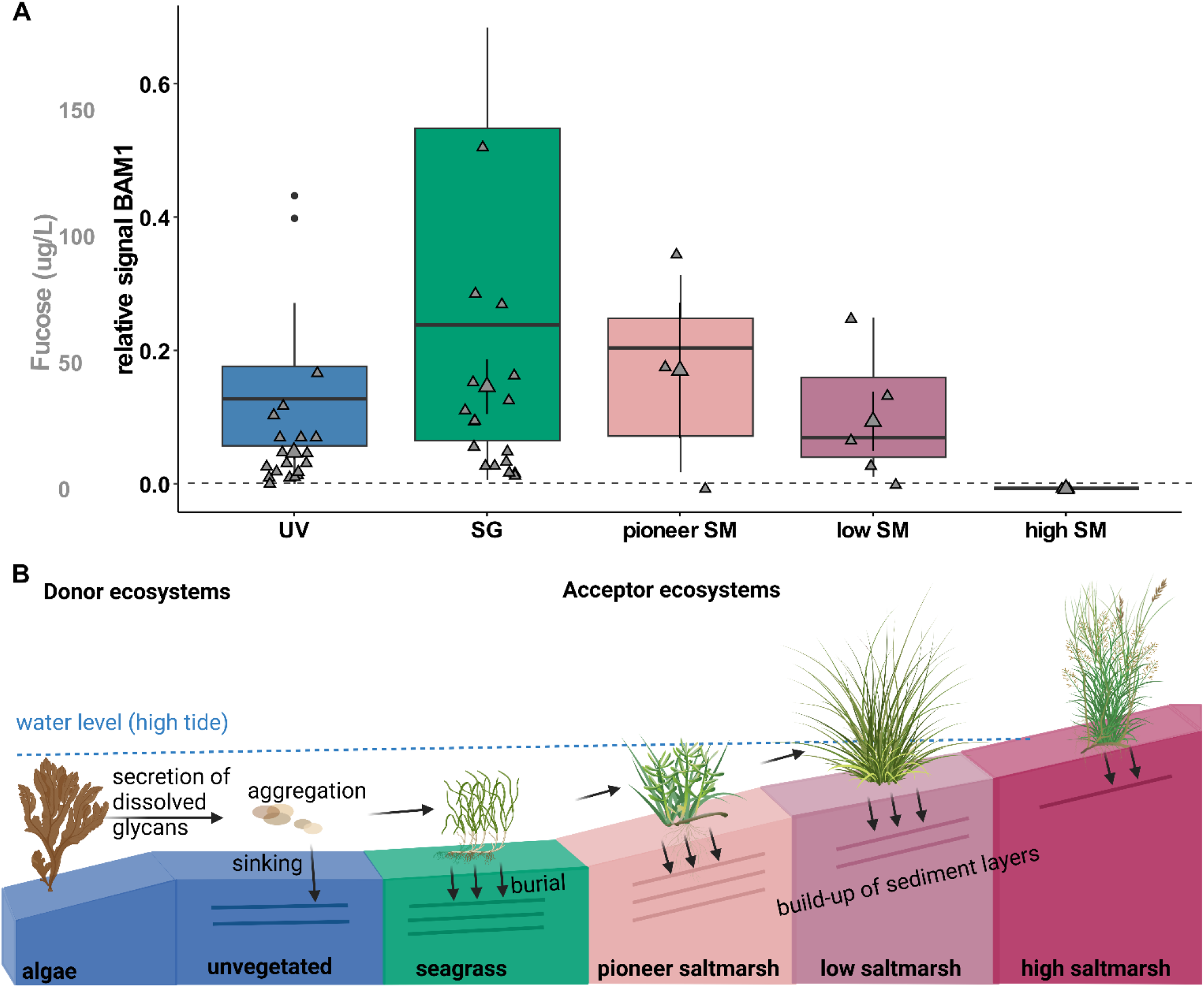
Algal glycan fucoidan is stored in coastal vegetated ecosystems. **A)** Relative signal for antibody BAM1 (indicating presence of fucoidan) binding to porewater extracts from unvegetated (UV; blue), seagrass (SG; green), pioneer, low and high (SM; rose to maroon). Relative signal is normalized to signal of MilliQ-water blank (dashed line). Mean fucose concentration in µg L^-1^ (mean ± s.e.m) for all ecosystems (grey). **B)** Schematic view on storage of algal-derived glycans in coastal vegetated ecosystems, using BioRender (Algal-Derived Carbon Transported into Coastal Vegetated Ecosystems. Created in BioRender, 2025). Donor ecosystems, such as algae secrete dissolved glycans, which may aggregate and be transported to coastal vegetated ecosystems. These acceptor ecosystems, reached by tidal waters may bury transported algal glycans and build-up of their sediment layers, accreting and storing their own and the carbon of donor ecosystems.

## 4 Discussion

By analysing the mono- and polysaccharide compositions of 93 sediment cores, we found consistent profiles across all samples, despite their origins in diverse coastal ecosystems ranging from tropical to temperate regions. This was surprising as we expected a major variation in those profiles due to the differing sources and processing pathways of DOC and POC associated with each ecosystem type. For example, up to half of mangrove net primary production is exported to the ocean as organic matter (Jennerjahn and Ittekkot, 2002) and seagrass is estimated to contribute to half of the surface sediment carbon pool of seagrass meadows (Kennedy et al., 2010). However, the observed similarity in relative monosaccharide abundances underneath all sampled ecosystems (**Fig. 1A**) suggests a glycan continuum, as previously described in surface waters (Aluwihare et al., 1997, 2002). The glycan continuum refers to the idea that structurally similar glycans persist after more labile organic matter has been broken down.

The diffusion map results highlight this consistency in glycan profiles across different ecosystems and locations (**Fig. 1B-C**). The major difference identified is driven by the relative glucose vs. fucose abundances in the samples, independent of the ecosystem and location. The fact that the relative abundance of fucose is low in high glucose samples and increases if glucose decreases indicates that fucose-rich oligosaccharides and polysaccharides might persist longer (Bligh et al., 2024; Miksch et al., 2024; Priest et al., 2023), while glucose rich glycans are broken down faster This hypothesis is supported by the findings that glucose-rich glycans such as laminarin are much easier to break down compared to fucose-rich glycans like fucoidan, which requires a myriad of enzymes to degrade (Becker et al., 2020; Sichert et al., 2020).

Remarkably, we detected a fucoidan antibody signal in the sediments across all coastal ecosystems. As fucoidan is predominantly produced and exuded by algal species (Koch et al., 2019), it can besides other algal-derived polysaccharides serve as a tracer for algal carbon sequestration and for tracing algal carbon from sources to sinks across ecosystems. Overall, polysaccharide screening of sediment samples revealed most binding to alginate, fucoidan and arabinogalactan (**Fig. S2, S3, Fig. 3**). Arabinogalactan-protein glycan and fucoidan are exuded by algal species, in particular by brown algae (Koch et al., 2019). Arabinogalactan-protein glycans can additionally stem from plant sources (Koch et al., 2019; Vidal-Melgosa et al., 2021). Fucoidan was shown to be actively and continuously produced and secreted in the mucilage layer of brown algae and diatoms (Buck-Wiese et al., 2023; Huang et al., 2021; Vidal-Melgosa et al., 2021). This continuous fucoidan secretion and the resulting increase in dissolved organic matter may subsequently promote an increasing formation of particulate organic matter. These particles can be exported with other carbon-containing compounds through the biological carbon pump (Engel et al., 2004; Iversen, 2023). The fucoidan and arabinogalactan-protein glycan signals measured here in coastal vegetated ecosystems, suggest their export to and storage in coastal sediments.

The consistent signal of algal-derived polysaccharides across the different ecosystems and locations and the higher abundance in coastal vegetated ecosystems reached by tidal waters compared to unvegetated areas and high saltmarsh areas (**Fig. 4**) highlights the importance of algae as donor ecosystems to blue carbon (Krause-Jensen et al., 2018; Krause-Jensen and Duarte, 2016). The root systems of seagrass meadows, saltmarsh plants and mangroves stabilize the sediment and the carbon stored within (e.g. Karimi et al., 2022). While around 7% of seagrass areas and up to 3% of saltmarsh and mangrove areas are lost annually (Mcleod et al., 2011), restoring and expanding these ecosystems could not only enhance local carbon storage but also increases the capacity to sequester carbon from distant algal sources, such as fucoidan.

In conclusion, coastal vegetated ecosystems store carbon dioxide partially as glycans, from which some have the potential to sequester carbon as a long-term storage. Especially the interaction of ecosystems makes it difficult to track carbon compounds back to their origin, resulting in difficulties budgeting their individual climate storage potential. Fucoidan might serve as a tracer for algal carbon stored in different locations. Overall, this study highlights the interplay and the interconnectedness of various ecosystems in storing large amounts of carbon. Together they function as a key-nature based solution for marine carbon dioxide removal.

## 5 Acknowledgments

The authors thank technicians at Max-Planck Institute for Marine Microbiology and Marum Centre for Environmental Research, University of Bremen, namely Tina Horstmann for assistance with carbohydrate microarray analysis, Katharina Föll for HPAEC-PAD measurements, as well as Theresa Fett and Mirco Wölfelschneider from Leibniz Centre for Tropical Marine Research (ZMT), Bremen, Germany for sampling of cores and processing of sediment cores. The authors thank Dariya Baiko and Michael Seidel from Institute for Chemistry and Biology of the Marine Environment (ICBM), Carl von Ossietzky Universität, Oldenburg and Jana Geuer and Tomasz Markowski from Max-Planck Institute for Marine Microbiology for sampling and processing of porewater. The authors thank José Ernesto Mancera-Pineda from Universidad Nacional de Colombia, Esteban Zarza González from Universidad del Sinú, Colombia, Jen Nie Lee from Faculty of Science and Marine Environment, Universiti Malaysia Terengganu and A. Aldrie Amir from Institute for Environment and Development (LESTARI), Universiti Kebangsaan Malaysia. Collections along the German North Sea coast in Lower Saxony and Schleswig-Holstein were made under the permit issued by the Lower Saxony Wadden Sea National Park Authority (Ref.no.: 01.2-22249-1-1.1 (60-8) / 2022) and the National Park Administration of the State Agency for Coastal Protection, National Park and Marine Conservation Schleswig-Holstein (Ref.no.: 3141-537.46) respectively. Collections along the German Baltic Sea coast near Massholm, Wendtorf, and Heiligenhafen were made under the permits issued by the Schleswig-Flensburg District Department for Environment Section for Nature Conservation, the Plön District Administrator, Lower Nature Conservation Authority, Office for the Environment (Ref.no.: 3106-3/127/0161), and the District Administrator Department Nature and Soil (Ref.no.: 6.20.2-3117-IV-22-Fr) respectively. Collections from Colombia along the peninsula of Barú were made under the permit from the Sub-Directorate for Management and Administration of Protected Areas (Ref.no.: 20222000120371). Collections from Malaysia were made under permit from the Ministry of Natural Resources, Environment and Climate Change (Access and benefit-sharing, Ref 974126). The permission to carry out research in Kilim Geoforest Park (Forest Reserve) in Kedah, Malaysia was obtained and approved by the Director-General of the Forestry Department of Peninsular Malaysia (Reference No.: JH/ 100 Jld. 33 (75)). The permit to enter Kilim Geoforest Park (Forest Reserve) in Kedah, Malaysia was obtained from the Langkawi District Forest Office (Permit No.: KL 72.2022). The authors acknowledge invaluable support from the Max-Planck-Society, the sea4SoCiety, CDRmare campaign in the German Marine Research Alliance.

## 6 Author contributions

I.H. and J.-H.H. designed research with contributions from all authors; I.H. performed sampling in Germany and total carbohydrate, monosaccharide quantification and ELISA antibody binding. A.A.M. performed microarray analysis. I.H., J.M. and J.-H.H. analyzed data and I.H. and J.H.H. led the manuscript production with contributions from all authors.

The authors declare no competing interest.

