## Supplementary material for "Fucoidan carbon is stored in coastal vegetated ecosystems"

### 1 Supplementary information

#### 1.1 Supplementary tables

**Table S1:** Sediment cores taken in Germany, Malaysia and Columbia in all four ecosystems: SG (seagrass), UV (unvegetated), SM (saltmarsh) and MG (mangrove). Information on sampling location, date and time.

| Country | Location | Ecosystem | Point | Longitude | Latitude | Date and time |
|---|---|---|---|---|---|---|
| Germany | NES | SG | P4 | 7.335864 | 53.686188 | 2022-08-16T08:43:00 |
| Germany | NES | SG | P5 | 7.335658 | 53.686212 | 2022-08-16T08:59:00 |
| Germany | NES | SG | P6 | 7.335419 | 53.686239 | 2022-08-16T09:13:00 |
| Germany | NES | UV | P4 | 7.336820 | 53.685264 | 2022-08-17T07:02:00 |
| Germany | NES | UV | P5 | 7.336531 | 53.685304 | 2022-08-17T07:13:00 |
| Germany | NES | UV | P6 | 7.336299 | 53.685328 | 2022-08-17T07:26:00 |
| Germany | NES | SM | P3 | 7.335585 | 53.684371 | 2022-08-15T10:46:00 |
| Germany | NES | SM | P6 | 7.335641 | 53.684276 | 2022-08-15T11:32:00 |
| Germany | NES | SM | P8 | 7.335455 | 53.683768 | 2022-08-15T13:59:00 |
| Germany | NES | SM | P9 | 7.335623 | 53.683454 | 2022-08-15T13:42:00 |
| Germany | HAH | SG | P4 | 8.823788 | 54.612684 | 2021-11-09T09:31:03 |
| Germany | HAH | SG | P6 | 8.823431 | 54.612663 | 2021-11-09T09:14:41 |
| Germany | HAH | SG | P8 | 8.822986 | 54.612680 | 2021-11-09T09:01:44 |
| Germany | HAH | UV | P4 | 8.811397 | 54.605695 | 2021-11-10T09:10:14 |
| Germany | HAH | UV | P5 | 8.811318 | 54.605757 | 2021-11-10T09:20:28 |
| Germany | HAH | UV | P6 | 8.811194 | 54.605855 | 2021-11-10T09:29:10 |

| | | | | | | |
|---|---|---|---|---|---|---|
| Germany | HAH | SM | P5 | 8.825151 | 54.604496 | 2021-11-08T14:20:59 |
| Germany | HAH | SM | P7 | 8.824193 | 54.605339 | 2021-11-08T13:55:06 |
| Germany | MET | SG | P4 | 8.967728 | 54.502967 | 2021-11-11T09:54:31 |
| Germany | MET | SG | P6 | 8.968376 | 54.502919 | 2021-11-11T09:41:19 |
| Germany | MET | UV | P6 | 8.964934 | 54.503152 | 2021-11-11T12:17:11 |
| Germany | MET | SM | P1 | 8.961645 | 54.503247 | 2021-11-10T15:01:18 |
| Germany | MET | SM | P4 | 8.961963 | 54.503353 | 2021-11-10T14:30:54 |
| Germany | SCH | SM | P4 | 10.024382 | 54.691559 | 2022-08-30T09:53:00 |
| Germany | SCH | SM | P5 | 10.025295 | 54.691736 | 2022-08-30T09:54:00 |
| Germany | SCH | SM | P6 | 10.026225 | 54.691997 | 2022-08-30T09:56:00 |
| Germany | SCH | SM | P7 | 10.023244 | 54.692271 | 2022-08-30T10:43:00 |
| Germany | WEN | SG | P5 | 10.287542 | 54.427274 | 2022-08-29T08:47:00 |
| Germany | WEN | SG | P7 | 10.286985 | 54.427172 | 2022-08-29T08:25:00 |
| Germany | WEN | SG | P9 | 10.286444 | 54.426960 | 2022-08-31T08:29:00 |
| Germany | WEN | UV | P3 | 10.281737 | 54.425132 | 2022-08-26T10:31:00 |
| Germany | WEN | UV | P5 | 10.281203 | 54.424871 | 2022-08-26T10:26:00 |
| Germany | WEN | UV | P7 | 10.280711 | 54.424579 | 2022-08-26T10:22:00 |
| Germany | WEN | SM | P5 | 10.290692 | 54.425844 | 2022-08-26T08:21:00 |
| Germany | WEN | SM | P7 | 10.291283 | 54.426592 | 2022-08-26T08:16:00 |
| Germany | WEN | SM | P9 | 10.291536 | 54.427105 | 2022-08-26T08:11:00 |
| Germany | HEI | SG | P3 | 11.024932 | 54.380240 | 2022-08-24T10:08:00 |
| Germany | HEI | SG | P6 | 11.024368 | 54.380322 | 2022-08-24T10:04:00 |
| Germany | HEI | SG | P8 | 11.024125 | 54.380358 | 2022-08-24T10:02:00 |
| Germany | HEI | UV | P3 | 11.023301 | 54.381132 | 2022-08-25T09:01:00 |
| Germany | HEI | UV | P5 | 11.022933 | 54.381358 | 2022-08-25T09:04:00 |
| Germany | HEI | UV | P7 | 11.022446 | 54.381428 | 2022-08-25T09:06:00 |
| Germany | HEI | SM | P1 | 11.021790 | 54.377104 | 2022-08-23T12:11:00 |

| | | | | | | |
|---|---|---|---|---|---|---|
| Germany | HEI | SM | P3 | 11.020723 | 54.379271 | 2022-08-23T13:16:00 |
| Germany | HEI | SM | P4 | 10.995120 | 54.377100 | 2022-08-24T07:49:00 |
| Germany | HEI | SM | P9 | 10.998305 | 54.378869 | 2022-08-24T07:31:00 |
| Malaysia | LAN | SG | P1 | 99.817006 | 6.447992 | 2022-09-29T00:26:00 |
| Malaysia | LAN | SG | P3 | 99.817083 | 6.448251 | 2022-09-29T00:23:00 |
| Malaysia | LAN | SG | P5 | 99.817260 | 6.448333 | 2022-09-29T00:19:00 |
| Malaysia | LAN | UV | P1 | 99.819578 | 6.449959 | 2022-09-29T23:49:00 |
| Malaysia | LAN | UV | P3 | 99.819896 | 6.450094 | 2022-09-29T23:54:00 |
| Malaysia | LAN | UV | P5 | 99.820262 | 6.450287 | 2022-09-29T23:59:00 |
| Malaysia | LAN | MG | P1 | 99.817696 | 6.448197 | 2022-09-24T02:47:00 |
| Malaysia | LAN | MG | P5 | 99.818622 | 6.448140 | 2022-09-24T06:56:00 |
| Malaysia | PAK | MG | P5 | 103.439177 | 4.660548 | 2022-10-11T04:51:00 |
| Malaysia | PAK | MG | P9 | 103.439555 | 4.660660 | 2022-10-11T04:35:00 |
| Malaysia | SET | SG | P1 | 102.740665 | 5.660896 | 2022-10-05T01:51:00 |
| Malaysia | SET | SG | P2 | 102.740617 | 5.660790 | 2022-10-05T01:54:00 |
| Malaysia | SET | SG | P4 | 102.740644 | 5.660663 | 2022-10-05T01:58:00 |
| Malaysia | SET | UV | P1 | 102.740820 | 5.660473 | 2022-10-05T03:17:00 |
| Malaysia | SET | UV | P2 | 102.740859 | 5.660461 | 2022-10-05T03:18:00 |
| Malaysia | SET | UV | P3 | 102.740893 | 5.660429 | 2022-10-05T03:20:00 |
| Malaysia | SET | MG | P2 | 102.739409 | 5.658089 | 2022-10-04T02:43:00 |
| Malaysia | SET | MG | P5 | 102.739269 | 5.657932 | 2022-10-04T02:58:00 |
| Malaysia | SET | MG | P8 | 102.739227 | 5.657740 | 2022-10-04T02:23:00 |
| Malaysia | SET | SM | P1 | 102.743881 | 5.658787 | 2022-10-09T03:01:00 |
| Malaysia | SET | SM | P3 | 102.744144 | 5.658643 | 2022-10-09T02:58:00 |
| Malaysia | SET | SM | P5 | 102.744319 | 5.658541 | 2022-10-09T02:55:00 |
| Columbia | 1 | SG | P3 | -75.681974 | 10.136862 | 2022-05-03T16:47:00 |
| Columbia | 1 | SG | P4 | -75.681995 | 10.136855 | 2022-05-03T16:48:00 |

| | | | | | | |
|---|---|---|---|---|---|---|
| Columbia | 1 | SG | P8 | -75.682115 | 10.137064 | 2022-05-03T16:53:00 |
| Columbia | 1 | UV | P2 | -75.682109 | 10.137113 | 2022-05-03T16:57:00 |
| Columbia | 1 | UV | P3 | -75.682165 | 10.137153 | 2022-05-03T16:56:00 |
| Columbia | 1 | UV | P4 | -75.682178 | 10.137119 | 2022-05-03T16:55:00 |
| Columbia | 1 | MG | P7 | -75.682718 | 10.136688 | 2022-05-01T14:44:00 |
| Columbia | 1 | MG | P8 | -75.682815 | 10.136645 | 2022-05-01T14:45:00 |
| Columbia | 1 | MG | P9 | -75.682996 | 10.136577 | 2022-05-01T14:46:00 |
| Columbia | 2 | SG | P3 | -75.630319 | 10.192668 | 2022-05-05T15:43:00 |
| Columbia | 2 | SG | P4 | -75.630338 | 10.192487 | 2022-05-05T15:39:00 |
| Columbia | 2 | SG | P5 | -75.630355 | 10.192527 | 2022-05-05T15:41:00 |
| Columbia | 2 | UV | P1 | -75.630572 | 10.192642 | 2022-05-07T15:54:00 |
| Columbia | 2 | UV | P2 | -75.630508 | 10.192592 | 2022-05-07T15:53:00 |
| Columbia | 2 | UV | P3 | -75.630435 | 10.192557 | 2022-05-07T15:52:00 |
| Columbia | 2 | MG | P6 | -75.630144 | 10.191658 | 2022-05-04T15:44:00 |
| Columbia | 2 | MG | P3 | -75.629953 | 10.191859 | 2022-05-04T15:29:00 |
| Columbia | 2 | MG | P2 | -75.630129 | 10.192157 | 2022-05-04T14:26:00 |
| Columbia | 3 | SG | P1 | -75.655608 | 10.171513 | 2022-05-08T21:21:00 |
| Columbia | 3 | SG | P6 | -75.655606 | 10.171255 | 2022-05-08T21:17:00 |
| Columbia | 3 | UV | P1 | -75.655774 | 10.171481 | 2022-05-09T14:50:00 |
| Columbia | 3 | UV | P2 | -75.655756 | 10.171415 | 2022-05-09T14:57:00 |
| Columbia | 3 | UV | P5 | -75.655592 | 10.171073 | 2022-05-09T14:56:00 |
| Columbia | 3 | MG | P5 | -75.655710 | 10.171036 | 2022-05-08T16:26:00 |
| Columbia | 3 | MG | P6 | -75.655767 | 10.170866 | 2022-05-08T16:21:00 |

**Table S2:** Porewater samples taken in North Sea, Germany: SG (seagrass), UV (unvegetated), SM (saltmarsh) and MG (mangrove). Information on sampling location, date and time.

| Country | Location | Ecosystem | Point | Longitude | Latitude | Date and time |
|---|---|---|---|---|---|---|

| | | | | | | |
|---|---|---|---|---|---|---|
| Germany | MET | UV | P8 | 8.965385406 | 54.50309346 | 2021-11-11T11:56:13 |
| Germany | MET | UV | P7 | 8.965154674 | 54.50312189 | 2021-11-11T12:05:02 |
| Germany | MET | UV | P6 | 8.964933863 | 54.50315194 | 2021-11-11T12:17:11 |
| Germany | MET | UV | P5 | 8.9646673 | 54.50318279 | 2021-11-11T12:28:43 |
| Germany | MET | UV | P4 | 8.964450012 | 54.50321822 | 2021-11-11T12:36:18 |
| Germany | MET | UV | P2 | 8.964039088 | 54.50325294 | 2021-11-11T13:01:43 |
| Germany | MET | UV | P1 | 8.963778231 | 54.50328101 | 2021-11-11T13:09:45 |
| Germany | MET | SG | P7 | 8.968679146 | 54.50289248 | 2021-11-11T09:35:04 |
| Germany | MET | SG | P6 | 8.968376408 | 54.50291868 | 2021-11-11T09:41:19 |
| Germany | MET | SG | P5 | 8.968031533 | 54.50295053 | 2021-11-11T09:47:01 |
| Germany | MET | SG | P3 | 8.967398385 | 54.50299082 | 2021-11-11T10:01:21 |
| Germany | MET | SG | P2 | 8.967011384 | 54.50301002 | 2021-11-11T10:08:13 |
| Germany | MET | SG | P1 | 8.966630373 | 54.50304336 | 2021-11-11T10:14:44 |
| Germany | MET | SM | P9 | 8.962556616 | 54.50367524 | 2021-11-10T13:29:20 |
| Germany | MET | SM | P7 | 8.962272025 | 54.50353863 | 2021-11-10T13:55:11 |
| Germany | MET | SM | P4 | 8.961963113 | 54.50335311 | 2021-11-10T14:30:54 |
| Germany | MET | SM | P3 | 8.961829112 | 54.50330163 | 2021-11-10T14:42:36 |
| Germany | NES | UV | P1 | 7.337581152 | 53.68513592 | 2022-08-17T06:15:00 |
| Germany | NES | UV | P2 | 7.337331868 | 53.68518217 | 2022-08-17T06:33:00 |
| Germany | NES | UV | P3 | 7.337111836 | 53.68522585 | 2022-08-17T06:48:00 |
| Germany | NES | UV | P4 | 7.336819655 | 53.68526396 | 2022-08-17T07:02:00 |

| | | | | | | |
|---|---|---|---|---|---|---|
| Germany | NES | UV | P5 | 7.336531409 | 53.68530355 | 2022-08-17T07:13:00 |
| Germany | NES | UV | P6 | 7.336299366 | 53.68532768 | 2022-08-17T07:26:00 |
| Germany | NES | UV | P7 | 7.336025374 | 53.6853751 | 2022-08-17T07:39:00 |
| Germany | NES | UV | P9 | 7.335573659 | 53.68543607 | 2022-08-17T07:50:00 |
| Germany | NES | UV | P8 | 7.335798533 | 53.68540797 | 2022-08-17T08:03:00 |
| Germany | NES | SG | P1 | 7.336522271 | 53.68607174 | 2022-08-16T07:34:00 |
| Germany | NES | SG | P2 | 7.336249798 | 53.68612155 | 2022-08-16T07:49:00 |
| Germany | NES | SG | P3 | 7.336034738 | 53.68615325 | 2022-08-16T08:26:00 |
| Germany | NES | SG | P4 | 7.335864038 | 53.68618797 | 2022-08-16T08:43:00 |
| Germany | NES | SG | P5 | 7.335658195 | 53.68621189 | 2022-08-16T08:59:00 |
| Germany | NES | SG | P6 | 7.335419068 | 53.68623867 | 2022-08-16T09:13:00 |
| Germany | NES | SG | P7 | 7.335206187 | 53.68625372 | 2022-08-16T09:27:00 |
| Germany | NES | SG | P8 | 7.335018737 | 53.68627393 | 2022-08-16T09:45:00 |
| Germany | NES | SG | P9 | 7.334821413 | 53.68629682 | 2022-08-16T09:57:00 |
| Germany | NES | SM | P1 | 7.334766191 | 53.6843031 | 2022-08-15T10:08:00 |
| Germany | NES | SM | P2 | 7.335256734 | 53.68429143 | 2022-08-15T10:34:00 |
| Germany | NES | SM | P5 | 7.335258637 | 53.68421474 | 2022-08-15T10:59:00 |
| Germany | NES | SM | P6 | 7.335641285 | 53.68427579 | 2022-08-15T11:32:00 |

### 1.2 Supplementary figures

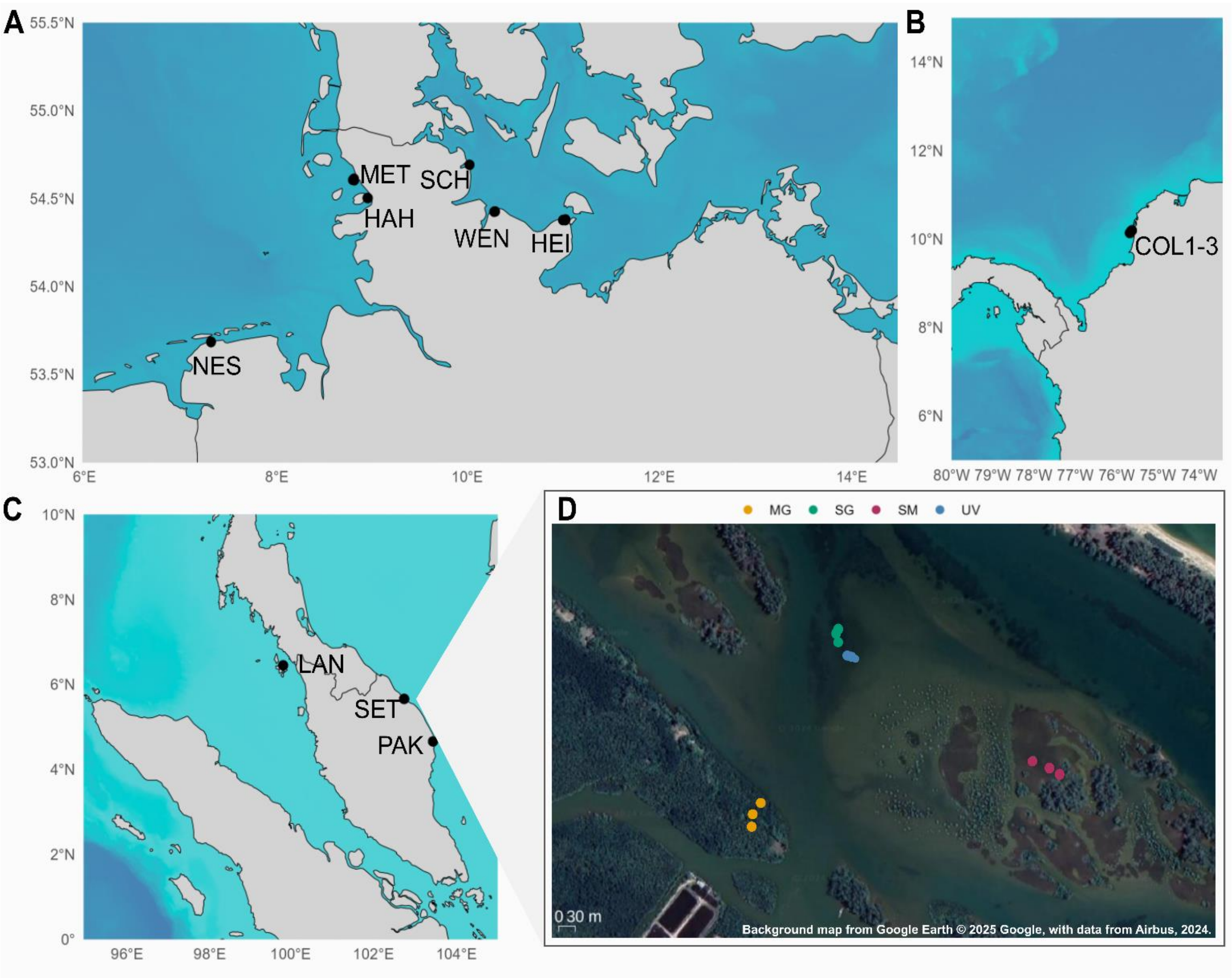


**Figure S1: Global sampling map of sediment cores of coastal vegetated ecosystems.** Bathymetrie is increasing with darker blue scale, bathymetry data is from NOAA (2022), plotted with R packages 'marmap' (Pante and Simon-Bouhet, 2013) and 'sf' (Bivand, 2021). **A)** Sampling locations along the North and Baltic Sea, Germany. NES: Nessmersiel, HAH: Hamburger Hallig, MET: Mettgrund, SCH: Schleimünde, WEN: Wendtorf, HEI: Heiligenhafen, **B)** in Columbia: COL sites 1 to 3 and **C)** in Malaysia, LAN: Langkawi, SET: Setiu, PAK: Paka. **D)** Example transect map, plotted with QGIS 3.22, background map from Google Earth © 2025 Google, with data from Airbus, imagery date 2024, of analysed transect point in ecosystems sampled in Setiu, Malaysia: mangroves (yellow), seagrass (green), saltmarsh (maroon), unvegetated (blue). Further information on analysed sampling points with locations are shown in supplementary **table S1**.

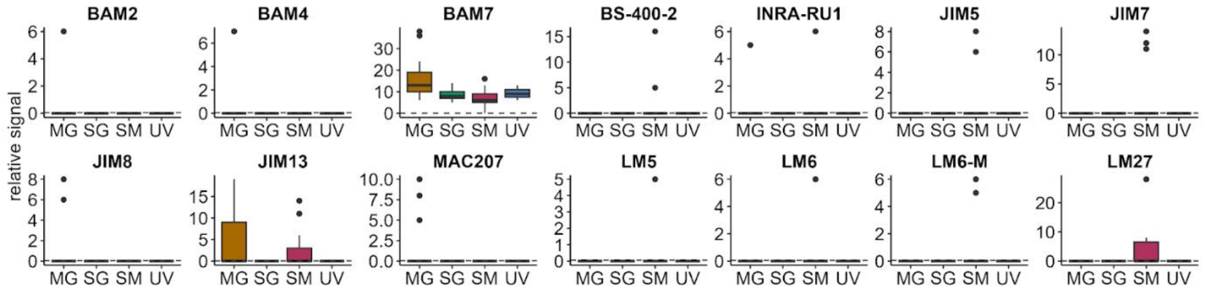


**Figure S2: Microarray antibody binding revealed presence of alginate, fucoidan, pectin, grass xylan and arabinogalactan.** Binding to BAM2 and BAM4 (sulfated epitope in sulfated fucan), BAM7 (alginate -pectin or fucoidan), BS400-2 ((1→3)-β-D-glucan), INRA-RU1 (rhamnogalacturonan I backbone), JIM5 (partially methyl-esterified/de-esterified HG), JIM7 (methyl-esterified HG), JIM8, JIM13 and MAC207 (all arabinogalactan protein glycan), LM5 ((1→4)-β-D-galactan), LM6 ((1→5)-α-L-arabinan), LM6-M ((1→5)-α-L-arabinan), LM27 (grass-xylan).

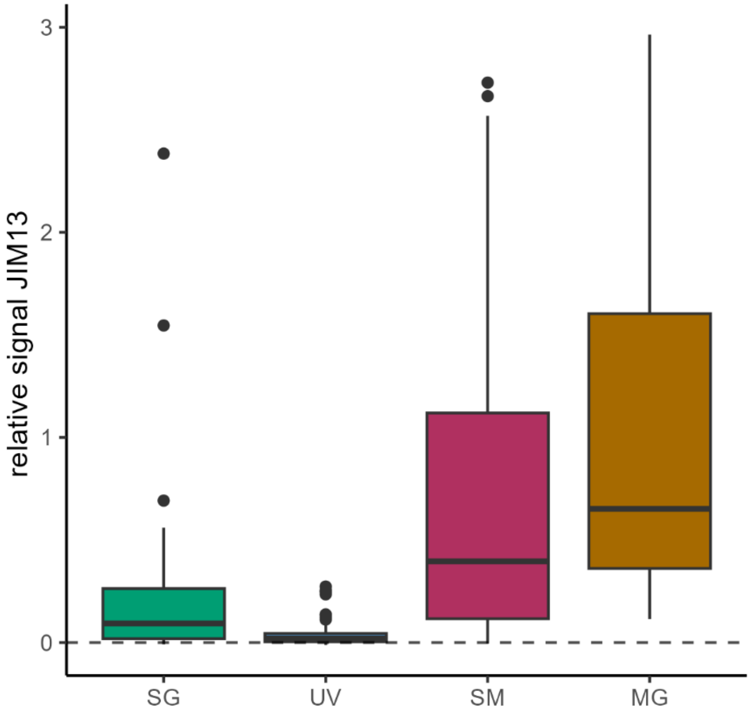

**Figure S3:** Relative antibody signal of JIM13 normalized to MilliQ-blank, representing arabinogalactans using ELISA method. Arabinogalactans (JIM13) found in saltmarsh (SM), mangrove (MG) and seagrass (SG) areas. No signal in unvegetated areas (UV).
